# A constellation of dysfunctional hybrid phenotypes enforces reproductive isolation between *Caenorhabditis* nematode species

**DOI:** 10.1101/2025.02.19.639108

**Authors:** Maia N. Dall’Acqua, Amanda L. Peake, Jalina Bielaska Da Silva, Dina Issakova, Asher D. Cutter

**Affiliations:** Department of Ecology & Evolutionary Biology, University of Toronto, Toronto, Ontario, M5S3B2 Canada

**Keywords:** Speciation, Reproductive isolation, Development, Hybridization, Nematodes, Caenorhabditis

## Abstract

The evolution of complete reproductive isolation hinges on the cumulative action of reproductive isolating barriers that can manifest throughout the life cycle of an organism. Consequently, a comprehensive understanding of the features underlying the origin and maintenance of species requires assessing the relative contributions of distinct barriers to overall reproductive isolation. Here we characterize multiple interrelated isolating barriers across various developmental stages of nematode sister species *Caenorhabditis remanei* and *Caenorhabditis latens*. We quantified F1 hybrid male sterility and characterized multiple phenotypic causes associated with developmental abnormalities in the germline as well as non-germline gonad and somatic tissues, uncovering a complex suite of developmental defects contributing to strong postzygotic reproductive isolation. Despite these multifarious isolating barrier traits, assays testing for interspecies sperm transfer under “choice” conditions did not yield evidence of premating isolation. In contrast to other *Caenorhabditis* species pairs, we also found no evidence that ectopic sperm migration acts as a postmating-prezygotic barrier. The constellation of phenotypic defects in hybrids points to a polygenic or highly pleiotropic basis for hybrid dysfunction and implicates more rapid evolution of intrinsic postzygotic reproductive isolation than prezygotic isolation in these organisms.

## Introduction

The origin, divergence and maintenance of species is contingent on the strength of reproductive isolation (Mayr 1942; Butlin and Stankowski 2020). A combination of pre- and postzygotic reproductive isolating barriers mediate the position of diverging populations along the speciation continuum, and can evolve to grow stronger to eventually prevent any genetic exchange through intermediary hybrid offspring (Coyne et al. 2004; Butlin and Stankowski 2020; Matute and Cooper 2021; Stankowski and Ravinet 2021). Therefore it is essential to detail both pre- and postzygotic barriers to characterize the mechanistic and genetic interactions underlying reproductive isolation and to assess their relative influence on the capacity of incipient species for continued divergence versus merger (Coyne et al. 2004; Roux et al. 2016). Incompletely isolated species, found in many taxa, point to diverse ways in which reproductive barriers act through distinct traits and at different life stages, highlighting the complex and sometimes-transient nature of the mechanisms that can underlie speciation (Ramsey et al. 2003; Sobel et al. 2010; Rometsch et al. 2020; Jiménez-López et al. 2023). A common assumption holds that prezygotic isolation exceeds postzygotic isolation in the early stages of divergence, especially for sympatric species, as prezygotic barriers limit the production of low-fitness hybrid offspring among individuals occupying the same area (Coyne and Orr 1989; Christie et al. 2022; Shaw et al. 2024). However, a variety of cases document strong postzygotic isolation largely in the absence of prezygotic barriers (Christianson et al. 2005; Bracewell et al. 2011; Brekke and Good 2014; Gustafsson et al. 2014). This discrepancy highlights the need to characterize the timing and relative influence of distinct isolating barriers in diverse organisms to more fully identify generalities about the underlying evolutionary processes that govern speciation. Through such analysis, we can address the outstanding questions about the complexity, correlation and predictability of pre- and postzygotic barriers that effectively restrict hybridization across developmental stages.

Prezygotic barriers that manifest before mating, premating isolation, can involve ecology-dependent (extrinsic) and ecology-independent (intrinsic) contexts and behaviors that inhibit the likelihood of successful mating. Niche differentiation (Schluter 2000; Nosil 2012), for example, between different *Rhagoletis* fly species confers increased receptivity to fruit odor from their respective host plants, which greatly reduces the chance of reproductive interactions between species (Linn et al. 2004). Ecologically-independent divergence in sexual signaling also can prevent mating should interspecies encounters occur, as for many female birds that tend to prefer conspecific male songs and plumage (Uy et al. 2018). Even with attempted mating, mechanical incompatibilities can contribute to prezygotic isolation, as in *Ohomopterus* ground beetles, where differences in body size and genital morphology prevent successful copulation between certain species (Nagata et al. 2007). Prezygotic reproductive isolation also can operate after mating but before fertilization due to postmating-prezygotic (PMPZ) barriers acting at the gametic level. For example, cryptic female choice can impede heterospecific fertilization through conspecific pollen or sperm precedence in some plants and animals (Price 1997; Geyer and Palumbi 2005; Howard et al. 2009; Montgomery et al. 2010). *Drosophila mojavensis* and *Drosophila arizonae* display another type of PMPZ barrier, such that female *D. mojavensis* are unable to effectively degrade the insemination reaction mass induced by *D. arizonae* males, impeding hybrid zygote formation (Kelleher and Markow 2007).

When prezygotic barriers do not prevent fertilization entirely, hybrid offspring may display postzygotic developmental dysfunction. In the extreme, this developmental dysfunction can manifest as inviability at multiple developmental stages or as sterility of adults. Such postzygotic reproductive isolation mediated by Dobzhansky-Muller incompatibilities (DMIs) is expected to accumulate over evolutionary time along with genomic divergence, as a result of negative effects of epistatic genetic interactions between divergent genes and/or gene regulatory elements (Orr 1995; Orr and Turelli 2001; Coyne et al. 2004; Mack and Nachman 2017; Simon et al. 2018; Cutter and Bundus 2020; Schneemann et al. 2024). Developmental irregularities can present at one or more life stages within hybrid individuals and a given DMI may affect one or more distinct cell types, tissues, or organs if they interface with a shared genetic interaction through pleiotropy (Cutter 2023). Hybrid sterility can evolve rapidly in males (Coyne and Orr 1989; Wu 1992; Wu et al. 1996; Christianson et al. 2005; Presgraves and Meiklejohn 2021), and speciation becomes irreversible when all hybrid offspring are infertile or inviable (Muller 1942; Kulmuni et al. 2020). Asymmetries between the sexes (Haldane’s Rule) and between reciprocal crosses (Darwin’s Corollary to Haldane’s Rule) in the degree of hybrid sterility or inviability occur commonly, often due to DMIs involving uniparentally inherited elements (Coyne 1985; Turelli and Moyle 2007; Cowell 2022; Cutter 2024).

Given the broad span of developmental stages that can be impacted by reproductive isolation in distinct or genetically correlated ways, it is essential to quantify the suite of traits that contribute to premating, PMPZ, and postzygotic reproductive isolation (Ramsey et al. 2003; Sobel et al. 2010; Rometsch et al. 2020; Jiménez-López et al. 2023). To do so, we must establish which stages in an organism’s life cycle are most susceptible to incompatibility, which traits are most receptive to evolutionary change, and how this change occurs. From these foundations, we may then begin to develop clear hypotheses about the role of pleiotropy in DMI effects across development, the genetic complexity of incompatibility in terms of potential for multi-locus DMIs or multiple distinct DMIs, and the potential for predictability in the evolution of intrinsic reproductive isolating mechanisms. To address these issues, it is helpful to exploit study systems that are amenable to manipulative mechanistic investigation. Here, we take advantage of *Caenorhabditis* nematodes as an emerging model for speciation research (Baird et al. 1992; Baird and Yen 2000; Woodruff et al. 2010; Dey et al. 2014; Cutter 2018; Bi et al. 2019).

Investigations of speciation with *Caenorhabditis* nematodes have accelerated since the discovery of sister species with incomplete reproductive isolation (Woodruff et al. 2010; Dey et al. 2012). The small physical size, small genomes, rapid generation time, transparent cuticle, determinate cell lineage, and experimental manipulability of these animals make them unusually well-suited to mechanistic investigation (Corsi et al. 2015). Although reproductive barriers have been studied in lab environments for several species, little evidence exists of premating isolation in *Caenorhabditis*, as several species mate readily under “no-choice” conditions (Garcia et al. 2007; Dey et al. 2012, 2014; Bundus et al. 2018). However, strong PMPZ isolation resulting from ectopic sperm cell migration has been documented in certain species pairs, including for *C. elegans* and *C. nigoni* (Ting et al. 2014, 2018) as well as between *C. macrosperma* and *C. nouraguensis* (Schalkowski et al. 2024). Postzygotic isolation quantified for one gonochoristic species pair, *Caenorhabditis remanei* and *Caenorhabditis latens,* shows they are still capable of producing fertile hybrid progeny, despite having diverged approximately 5 million years ago (Dey et al. 2012). Previous research found significant embryonic inviability among F1 hybrid eggs (28.5% for *C. remanei* maternal, 33.4% for *C. latens* maternal), manifesting primarily during or after gastrulation. There is also a striking female sex bias in surviving F1 hybrid offspring from hybrid crosses with *C. remanei* as the maternal parent, where 74% of surviving progeny were female (Dey et al. 2012, 2014; Bundus et al. 2018). Hybrid F1 males from the reciprocal cross were all fertile, as were hybrid females from both cross directions, though F2 hybrid breakdown emerged from substantial embryonic arrest of F2 offspring (79.5% for *C. remanei* grandmaternal, 73.7% for *C. latens* grandmaternal) (Dey et al. 2014). Backcross analysis indicated that hybrid male sterility appears to have a simpler genetic basis than hybrid inviability and that postzygotic isolation involves a combination of X-autosome, autosome-autosome, and mito-nuclear DMIs (Bundus et al. 2018). Of the viable F1 hybrid males from the more severely-affected *C. remanei*-maternal crosses, 92-96% were sterile and 95% displayed abnormal gonad morphologies (Dey et al. 2014), although fertility and gonad morphology were not assessed in the same individuals simultaneously. Anecdotal observations of these hybrids highlighted other phenotypic traits that might also be associated with sterility (Dey et al. 2014), including male tail reproductive structures used for mate chemosensation, copulation, and insemination in *Caenorhabditis* (Liu and Sternberg 1995; Garcia et al. 2007; Ebert and Bargmann 2024).

In this study, we aim to inform general principles in the evolution of reproductive isolation by expanding our understanding of the phenotypic causes of pre- and postzygotic isolation between *C. remanei* and *C. latens*. To assess premating barriers, we quantified the incidence of successful sperm transfer amongst conspecific and heterospecific males and females under mutual “choice” conditions. Ectopic sperm migration was examined as a possible PMPZ barrier by measuring the location and quantity of ectopic sperm in the bodies of females after mating. To complement known postzygotic dysfunction phenotypes in embryogenesis, we investigated a suite of other traits involved in postzygotic isolation, including hybrid male fertility, testis development, spermatid morphology and number, and male tail reproductive structures. By comprehensively characterizing traits that contribute to reproductive barriers across the life cycle, we highlight the complex and multi-faceted nature of reproductive isolation and its underlying mechanisms.

## Methods

### Strains and maintenance

We maintained *C. remanei* (strains PB219 and VX3) and *C. latens* (strain VX88) according to standard *C. elegans* protocols (Stiernagle 2006). PB219 originates from a wild isolate collected in Dayton Ohio, USA (Baird 1999), VX3 was collected at the Koffler Scientific Reserve near Toronto ON, Canada (Dey et al. 2014) and VX88 originates from a wild isolate collected near Wuhan, China (Dey et al. 2012). All strains were maintained at 25°C on 6cm diameter petri dishes containing 2% Nematode Growth Media (NGM)-agar seeded with *Escherichia coli* strain OP50 as a food source. We removed bacterial and fungal contaminants from the populated petri dishes through standard bleaching protocols when necessary prior to performing experimental assays (Stiernagle 2006).

### Assortative mating of species

To assess the propensity for interspecies mating in arenas containing both sexes of both *C. remanei* (strain VX3) and *C. latens* (strain VX88), we quantified conspecific and heterospecific sperm transfer. 24 hours prior to experimental testing, ∼75 young adult males of each species were immersed in 300μL M9 containing 5μL of 50mM MitoTracker Red CMXRos, a fluorescent vital dye that allows for the visualization of spermatozoa (Kubagawa et al. 2006; Ting et al. 2014, 2018a). 2.5 hours prior, ∼75 unmated young adult females of each species were immersed in 51.5μL of 50uM CellTracker Green BODIPY, another fluorescent vital dye, to facilitate visual species identification. We then established “choice” mating arenas with a mixed population of 10 males and 10 females of each species (40 animals total per petri dish) with a single *E. coli* spot in the centre: 3 replicate 3cm-diameter plates containing MitoTracker Red-dyed *C. remanei* males (unstained *C. remanei* females) and CellTracker Green-dyed *C. latens* females (unstained *C. latens* males) and another 3 replicate plates of worms with reciprocal staining. We also set up comparable conspecific-only mating plates in triplicate for each species as controls, each containing 20 females and 20 males of the same species with half of the males dyed with MitoTracker Red and half of the females dyed with CellTracker Green. Plates were placed in a light-proof box to prevent photodegradation of the fluorescent dyes and incubated at 25°C for 4 hours to allow for mating. Females were then transferred to 5% agarose pads on microscope slides to prevent damage during coverslip application and immobilized in 10uL of 50mM NaN_3_. In some cases, 1-2 females died or crawled off the plate and were not scored. Slides were examined at 20X and 40X magnification using an Olympus BX51 microscope and Leica EL6000 light source and the appropriate fluorescence filters to score the species identity of each female and the presence or absence of fluorescent sperm in their reproductive tracts.

Statistical analyses were performed in R v4.4.1 (R Core Team 2024), and all data and scripts are available on Zenodo (DOI: <u>10.5281/zenodo.14888936</u>) and GitHub (github.com/Cutterlab/Constellation_of_RIs). We used a mixed effects logistic regression model to evaluate the effect of cross direction on the frequency of successful sperm transfer. Cross direction was included as a fixed effect and experimental block as a random effect. Species identity of females dyed with CellTracker Green and replicate mating plate accounted for negligible variance in sperm transfer success and were excluded from the model. We then calculated pairwise contrasts of estimated marginal means for all combinations of conspecific and heterospecific crosses to compare sperm transfer frequency between cross directions. Post hoc Tukey tests were applied to correct for multiple comparisons. To test for baseline differences in sperm transfer frequency of pure species crosses, we ran a mixed effects logistic regression model which assessed the frequency of successful sperm transfer on conspecific-only mating plates with replicate mating plate as a random effect nested within species identity.

### Ectopic migration of sperm cells

To investigate whether heterospecific sperm can migrate ectopically in females, as occurs following heterospecific mating of some *Caenorhabditis* species (Ting et al. 2014, 2018a; Schalkowski et al. 2024), we conducted “no-choice” matings between *C. remanei* (strain VX3) and *C. latens* (strain VX88) and then visualized the location of sperm within the females’ bodies. Unmated adult males were dyed with MitoTracker Red CMXRos 24 hours ahead of the assay to visualize transferred sperm. 6 replicate plates each containing 6 unmated adult females and 12 unmated adult males were set up for both conspecific and reciprocal heterospecific cross directions. After 2 hours of mating opportunity in a light-proof box at 25°C, we removed the males and left the females for an additional 2 hours to allow for possible ectopic sperm migration. The females were then mounted on microscope slides as described for the assortative mating experiments and scored for fluorescent sperm at 40X magnification. We quantified the presence, location (uterus and spermathecae; gonad; somatic body cavity), and quantity of sperm and severity of ectopic sperm migration in females using simplified criteria based on Ting et al. 2018 (Table 1).

**Table 1.** Incidence, location and severity of ectopic sperm migration observed in *C. remanei* and *C. latens* females after conspecific and heterospecific crosses.

| Cross (♀x♂) | n | #♀ with ectopic sperm** | Sperm location |  |  | Severity* |  |  |
| --- | --- | --- | --- | --- | --- | --- | --- | --- |
|  |  |  | Uterus | Gonad | Soma | Mild | Moderate | Severe |
| *C. remanei* x *C. remanei* | 38 | 5 | 38 | 4 | 3 | 2 | 2 | 1 |
| *C. remanei* x *C. latens* | 35 | 7 | 35 | 7 | 1 | 2 | 4 | 1 |
| *C. latens* x *C. remanei* | 35 | 9 | 35 | 9 | 2 | 3 | 4 | 2 |
| *C. latens* x *C. latens* | 32 | 12 | 32 | 12 | 5 | 8 | 2 | 2 |

\*Severity was scored as mild (~1-10 ectopic sperm), moderate (~11-50 ectopic sperm), and severe (~51+ ectopic sperm) irrespective of ectopic sperm location.
\*\*Ectopic sperm was identified as sperm found in any tissue other than the uterus.

We used a mixed effects logistic regression model in R to investigate the influence of cross direction on frequency of ectopic sperm migration in females following mating. Replicate mating plate was included as a random effect. We then computed estimated marginal means to generate pairwise contrasts for all possible combinations of cross direction and applied post hoc Tukey tests to correct for multiple comparisons. Fisher’s exact tests evaluated differences in the location and severity of ectopic sperm migration between cross types.

### Fertility, gonad morphology and presence/absence of spermatids in F1 hybrid males

We separated unmated *C. remanei* (PB219) and *C. latens* (VX88) L4 males and females from age-synchronized petri dishes 24 hours in advance of crosses to prevent conspecific fertilization of females. We then incubated 7 replicate mating plates of 4 young adult *C. remanei* females and 4 young adult *C. latens* males at 25°C for 24 hours, after which all females were transferred to freshly seeded 6cm plates to lay eggs. Females remained on these plates for 48 hours while F1 offspring matured. Young adult F1 hybrid males were transferred individually to separate 6cm seeded plates containing 3 unmated adult *C. remanei* females and 3 unmated adult *C. latens* females. We monitored these plates every 24 hours to check for the presence of copulatory plugs, new eggs, offspring, and arrested embryos. Males on plates that contained no eggs or offspring within 2 days were categorized as sterile.

All fertile F1 hybrid males and a subset of infertile hybrid males were removed from mating plates after scoring and prepared for microscopy. A subset of hybrid males were prepared for only DIC imaging (n=10) and the remaining males (n=47) were DAPI stained and imaged. Individuals that were only DIC imaged were mounted on microscope slides containing 5% agarose pads and examined with a differential interference contrast (DIC) filter at 20X magnification and imaged using CellSens software (RRID: SCR_014551; Olympus corp). We scored gonads as either “normal” or “abnormal”, where abnormal gonads showed noticeable deviations from pure species morphology. Abnormal gonads were further classified into “mild” and “severe”, with mild defects showing only slight deviation from typical morphology, such as a slight anterior or posterior shift in gonad position or incomplete migration of the gonad arm. Gonads comprised of clumped tissue that did not resemble a normal gonad structure were classified as severe.

Worms that were prepared for both DAPI and DIC imaging were washed in PBS and fixed in −20°C methanol for 10 minutes before being DAPI stained. After 5 minutes, the worms were then resuspended in 15μL of 75% glycerol before being placed atop 5% agarose pads on new microscope slides. The condensed nuclei of DAPI-stained spermatids form a distinct clustering pattern in the proximal gonad, allowing for visualization under fluorescence. Each worm was scored for presence or absence of spermatids, and those containing spermatids were classified as having a “normal” or “excessive” number of spermatids compared to pure species males.

### Spermatid morphology

We extracted spermatids from both parental species as well as F1 hybrid males produced from reciprocal heterospecific crosses. To generate F1 males for both hybrids and parental species, 5 males of a given species with 5 unmated females of either the same or the other species (*C. remanei* strain PB219 and *C. latens* strain VX88) on one of 4 separate 6cm plates seeded with *E. coli* and incubated at 25°C for 24 hours. We then transferred mated females to individual 3.5cm plates for egg laying. Since any mating attempts between males and females on the plate would result in the loss of spermatids present in the males, we isolated all imaged males at the L4 stage onto a separate plate with no females present.

Each male was placed on a glass microscope slide in 10μL of sperm medium (50mM HEPES, 25mM KCl, 45mM NaCl, 1mM MgSO4, 5mM CaCl2, 10mM Dextrose) (Singaravelu et al. 2011) to ensure that spermatids retained their size and shape upon extraction. A 25G needle was used to cut the male just below the pharynx to allow for spermatid release from the body. The slides were then visualized at 100x magnification using oil immersion microscopy and imaged using CellSens software (RRID: SCR_014551; Olympus corp). Spermatid cross-sectional area (in square pixels) was measured using CellProfiler (Carpenter et al. 2006) and ImageJ v1.5 (Schneider et al. 2012), which gave comparable results. Additionally, ImageJ was used to quantify spermatid roundness as (4*π*area)/(perimeter^2^) and spermatid circularity as (4*area)/(π*(major axis length)^2^). We dissected a total of 20 individuals per cross type, and F1 hybrid males that did not produce spermatids were excluded from analysis. Generalized linear mixed models with parental cross as a fixed effect and F1 hybrid male as a random effect were used to evaluate the influence of parental cross on spermatid area, circularity, and roundness. Pairwise contrasts of estimated marginal means were evaluated for all combinations of conspecific and heterospecific crosses for each model followed by post hoc Tukey tests to adjust for multiple comparisons.

### Number of Spermatids

We generated F1 males for both parental species and F1 hybrid males from both cross directions as previously described in the “Spermatid morphology” section. Males were then DAPI stained and prepared for imaging as previously described in the “Fertility, gonad morphology, and presence/absence of spermatids in F1 hybrid males” section. We examined and imaged F1 males using an Olympus BX51 microscope and CellSens software (RRID: SCR_014551; Olympus corp) at 40X magnification. We took images at multiple depths of view where each image had an overlapping set of spermatids within focus so that we could account for all spermatids in each male. We used GIMP (The GIMP Development Team 2019) to create an overlay for each worm where we marked the location of all DAPI stained spermatid nuclei over all images for a given male. Overlays were then imported into DotCount v1.2 (Reuter 2012) software to count the total number of marked spermatids for each male. A total of 20 individuals were dissected per cross type, and F1 hybrid males that did not produce spermatids were excluded from analysis. We used a negative binomial generalized linear model to determine the effect of parental cross on spermatid number, and computed estimated marginal means to conduct pairwise comparisons between parental crosses. Post hoc Tukey tests were used to correct for multiple comparisons.

### Tail morphology of hybrid male reproductive structures

To characterize and quantify defects in external reproductive structures of F1 hybrid males, we conducted controlled crosses within and between *C. remanei* (strain PB219) and *C. latens* (strain VX88). Tails of the resulting F1 hybrid and pure species males were visualized at 40x using DIC microscopy. We set up 8 batches of reciprocal heterospecific crosses and 5 batches of conspecific crosses consisting of 5 replicate plates of 2 unmated young-adult males and 4 unmated young-adult females and incubated them at 25°C for 16-20 hours. After F1 offspring had matured for ∼48 hours, adult F1 males were removed from each plate, placed on microscope slides with 5% agarose pads, and visualized using DIC microscopy and imaged using CellSens software (RRID: SCR_014551; Olympus corp). We assessed each tail for the following abnormalities: fused rays, missing rays, deformed fan, abnormal tail shape, intact cuticle, asymmetric tail, or “other”. Tails were scored as “normal” if no defects were present and “abnormal” if one or more defects were observed.

We used a mixed effects logistic regression model with parental cross as a fixed effect and experimental block as a random effect to evaluate the influence of parental cross on the frequency of F1 hybrid male tail defects. Pairwise contrasts of estimated marginal means were computed to evaluate differences in tail defect frequency between all combinations of conspecific and heterospecific crosses. Tukey post hoc tests were applied to correct for multiple comparisons.

## Results

### No evidence of assortative mating by species

To determine whether prezygotic barriers exist between *C. remanei* and *C. latens*, we tested for behavioral species discrimination during mating. While males and females of these different species will readily mate with one another in “no choice” mating scenarios (Dey et al. 2012, 2014; Bundus et al. 2018), we assessed the degree of successful interspecies mating in “choice” mating arenas with both sexes of both species present, inferred from sperm transfer (Figure 1c) (Ting et al. 2014, 2018). We found that the incidence of successful sperm transfer was similar between heterospecific cross directions and *C. latens* conspecific matings, but we unexpectedly observed significantly fewer inseminated females from conspecific *C. remanei* matings than from heterospecific matings and marginally fewer inseminated females than from *C. latens* conspecific matings (Figure 1a). Interestingly, it appears that this reduced capacity of *C. remanei* individuals to mate conspecifically depends on the presence of *C. latens*, as we did not observe any differences in successful sperm transfer between plates containing only conspecific individuals and those containing both sexes of both species (Figure 1b).

**Figure 1.**
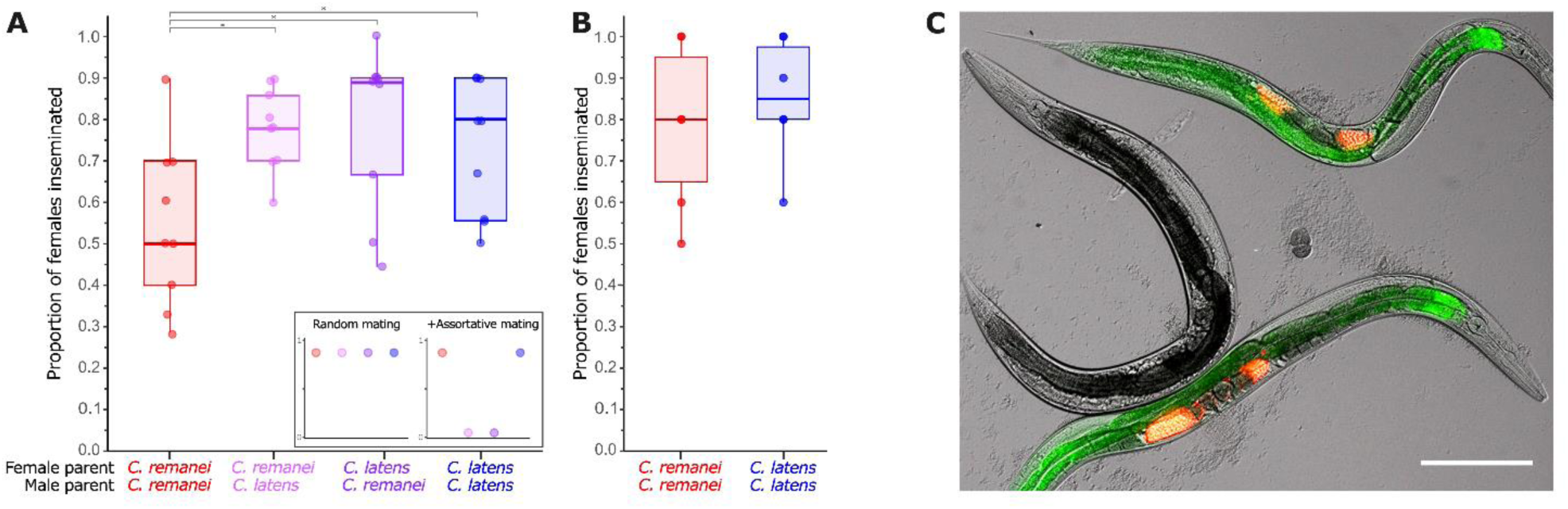
**(A)** Proportion of *C. remanei* and *C. latens* females inseminated by *C. remanei* and *C. latens* males under choice mating conditions. Points represent the proportion of females inseminated on one replicate mating plate containing 10 females of each species and 10 males of each species. Inset illustrates the expected patterns for random mating and positive assortative mating of the species. We observed no significant difference in the proportion of *C. remanei* and *C. latens* females inseminated through heterospecific matings versus *C. latens* conspecific matings (*P*>0.05 for all Tukey-corrected pairwise comparisons). Fewer females were inseminated in *C. remanei* conspecific crosses than in either heterospecific cross direction (*C. remanei* conspecific vs *C. remanei* maternal heterospecific: z=−3.012, Tukey-corrected *P*<0.05; *C. remanei* conspecific vs *C. latens* maternal heterospecific: z=−3.213, Tukey-corrected *P*<0.01) and in *C. latens* conspecific crosses, although this difference was only marginally significant (z=−2.443, Tukey-corrected *P*=0.0693). **(B)** Proportion of *C. remanei* and *C. latens* females inseminated on mating assay plates containing only conspecifics. The proportion of conspecifically inseminated *C. remanei* and *C. latens* females do not differ significantly (z=−0.03, *P*>0.05). **(C)** Image of two *C. remanei* females (top and bottom) and one *C. latens* female (center) after “choice” mating opportunity. Image is an overlay of DIC imaging as well as TRITC and FITC fluorescence filters. Red areas indicate *C. latens* sperm dyed with MitoTracker Red CMXRos and green areas signify CellTracker Green BODIPY in the digestive tract. Scale bar represents 100µm.

Overall, the similar or elevated likelihood of heterospecific matings compared to conspecific matings even under “choice” conditions indicates an absence of strong behavioral barriers to interspecies mating between *C. remanei* and *C. latens*, at least in a benign laboratory environment. Because we directly observed sperm cells in the reproductive tract of all mated females, it is also unlikely that mechanical barriers related to mating or sperm transfer contribute to pre-zygotic isolation.

### Incidence of ectopic sperm cell migration similar for heterospecific and conspecific crosses

Sexually antagonistic coevolution is predicted to have influenced the divergence of reproductive traits among *Caenorhabditis* species (Ting et al. 2014, 2018; Cutter et al. 2019). Mismatches between sperm competitiveness and female reproductive tract characteristics allow for the harmful ectopic migration of sperm cells following heterospecific matings between several combinations of *Caenorhabditis* species (Ting et al. 2014; Schalkowski et al. 2024). The prevalence of this phenomenon among some *Caenorhabditis* nematodes, and incomplete pre-mating isolation in *C. remanei* and *C. latens*, led us to investigate whether ectopic sperm cell migration, or “sperm invasion” in its extreme form, contributes a (PMPZ) isolating barrier between *C. remanei* and *C. latens*.

Fluorescence microscopy revealed that sperm frequently migrated ectopically after both heterospecific (26% of n=35 *C. latens* females; 20% of n=35 *C. remanei* females) and conspecific matings (34% of n=32*C. latens* females; 13% of n=38 *C. remanei* females) over the 4 h timespan of the assay (Table 1). However, we observed no significant difference between cross types in the proportion of females containing ectopic sperm (Tukey-corrected *P*>0.05 for 6 pairwise comparisons between crosses) (Table 1). We also observed no significant difference in migration severity among females from different cross types (Fisher’s exact test, *P*=0.342), with 55% of females across all cross types containing ectopic sperm being classified as moderate or severe cases (Table 1). Among females that experienced some degree of ectopic sperm cell migration (n=33), all but one individual contained sperm in the proximal gonad among the maturing oocytes, and 33% contained sperm that had migrated into the pseudocoelomic cavity (soma) and could be seen in regions near the head or tail (Table 1; Supplementary Figure 1). Such females often were dead by the time of imaging. Given that ectopic sperm has been shown to decrease reproductive output and lifespan in other *Caenorhabditis* species (Ting et al. 2014; Schalkowski et al. 2024), the relatively high degree of sperm invasiveness seen in this experiment likely compromises female reproductive capabilities and may represent a source of intraspecific sexual conflict rather than a source of interspecies reproductive isolation per se.

### Disrupted spermatogenesis partly explains hybrid male sterility as a result of defective gonadogenesis

We confirmed that most F1 hybrid males are sterile when they have *C. remanei* as the maternal parent (Dey et al. 2012, 2014; Bundus et al. 2018), observing that just 6 of 171 (3.5%) F1 hybrid males were fertile from *C. remanei* maternal heterospecific crosses. Distinct *C. remanei* maternal individuals of strain PB219 did not differ significantly in the incidence of fertility of their hybrid sons (Fisher’s exact test, *P*>0.05). Previous work also documented frequent gonadogenesis defects for this class of F1 hybrid males (Dey et al. 2014). To address whether defects in gonadogenesis still permit spermatogenesis, and whether disrupted gonadogenesis directly causes hybrid male sterility, we examined in more detail the gonad morphology for 61 of these *C. remanei-latens* F1 males, including the 6 that were fertile. We found that 9 of these males (15%) showed normal gonad morphology whereas 18 males, including 2 fertile males, displayed mild gonad abnormalities (30%), and 34 males (56%) displayed severe abnormalities (all of which were sterile) (Table 2). For a further subset of 43 sterile males, we assessed whether spermatids were present. We found that 7 of 16 sterile F1 males (43.8%) with mild gonad abnormalities did not contain spermatids, while 17 of 28 (60.1%) males with severely malformed gonads did not contain spermatids and 4 (14%) contained an abnormally excessive number of spermatids (Table 2). Thus, the capacity to produce sperm strongly associates with overall capacity to develop a gonad properly. Hybrid male gonad morphogenesis and sperm production contribute in a correlated manner to the incidence of hybrid male fertility versus sterility. Disrupted gonadogenesis nonetheless can sometimes permit successful spermatogenesis and, in some instances, leads to internal accumulation of spermatids, likely due to an inability to release them through the cloaca (see Discussion).

**Table 2.** Fertility, spermatid presence and gonad morphology in *C. remanei* (maternal) x *C. latens* (paternal) F1 hybrid males.

|  |  |  | Gonad morphology |  |  |
| --- | --- | --- | --- | --- | --- |
|  | n | Spermatids present | Normal | Mild defect | Severe defect* |
| Fertile | 6 | 6/6 | 4 | 2 | 0 |
| Infertile | 55 | 19/43 | 5 | 16 | 34 |

\*4 infertile individuals with severe gonad defects were also observed to have an excessive amount of sperm in their bodies relative to pure species males.

### Spermatid morphology is disrupted in interspecies hybrids

Given the capacity of at least some F1 males from both heterospecific cross directions to make spermatids, we tested for the potential of spermatid cells to manifest defects. We dissected F1 hybrid males from both cross directions to isolate spermatids and measured spermatid size, circularity, and roundness. Measures of spermatid size (cross-sectional area), circularity and roundness were lower in *C. remanei-latens* F1 hybrid males crosses compared to *C. remanei* and *C. latens* pure species males (Figure 2a-c). We also measured the number of spermatids present in F1 pure-species and hybrid individuals from a separate set of matings and found that *C. remanei-latens* F1 hybrid males showed a trend of fewer spermatids than *C. remanei* and *C. latens* males or F1 male hybrids from the reciprocal cross (*C. latens-remanei* F1 hybrid males), albeit not significant after multiple test correction (Figure 2d).

**Figure 2.**
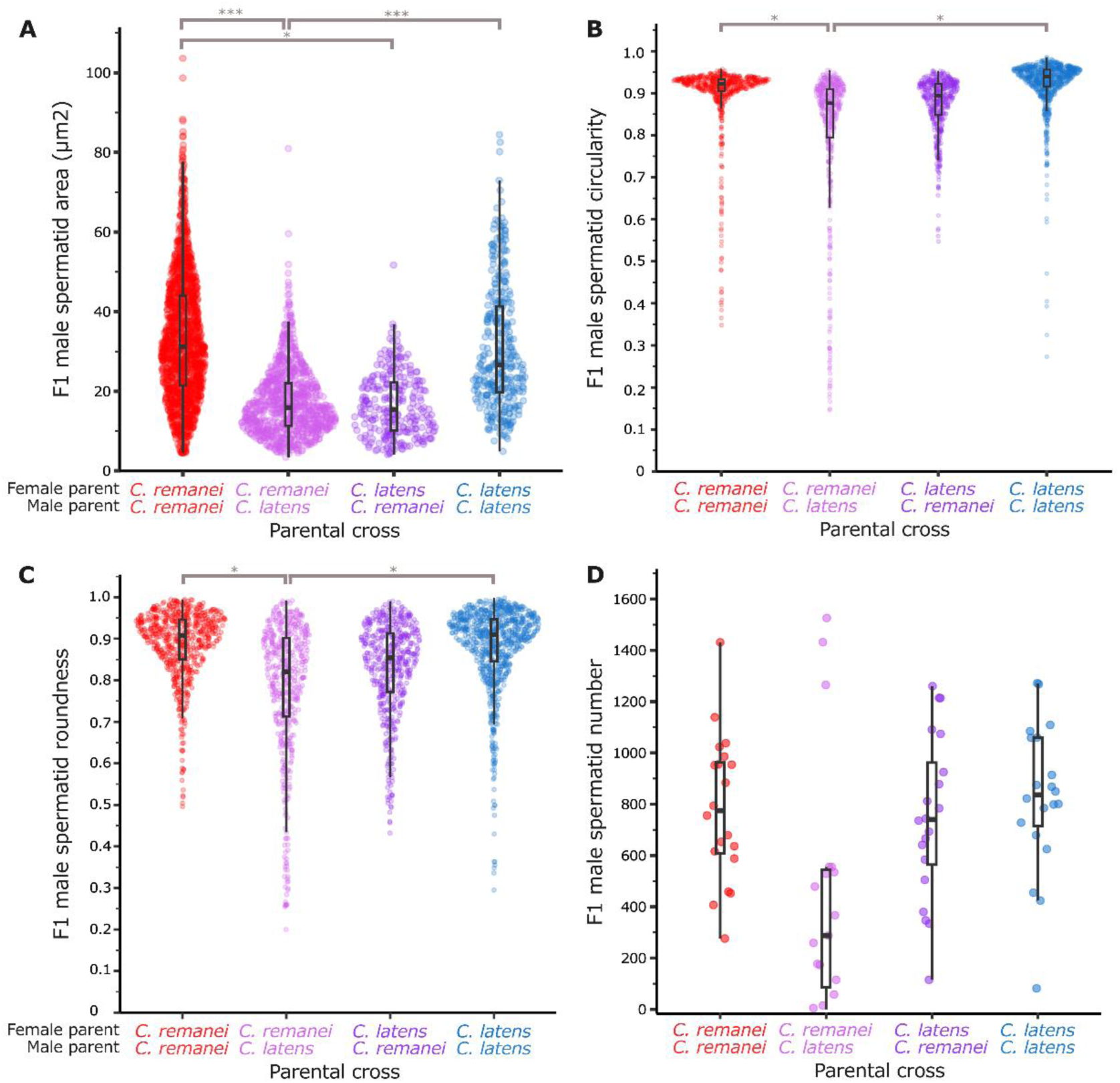
Quantification of spermatid phenotypes as a function of parental cross type. **(A)** Cross-sectional spermatid area is smaller in *C. remanei-latens* F1 hybrid males compared to *C. remanei* and *C. latens* pure species males (n_males_=22 and n_spermatids_=893 for *C. remanei-latens* F1 hybrid males, n_males_=25 and n_spermatids_=1444 for *C. remanei* pure species males, n_males_=12 and n_spermatids_=342 for *C. latens* pure species males; z=−6.361, Tukey-corrected *P*<0.0001; z=4.648, *P*<0.0001 respectively), and smaller in *C. latens-remanei* F1 hybrid males compared to *C. remanei* pure species males (n_males_=9 and n_spermatids_=269 for *C. latens-remanei* F1 hybrid males; z =−3.139, Tukey-corrected *P*<0.01). **(B)** *C. remanei-latens* F1 hybrid males have spermatids with lower circularity (after inverse transformation) than *C. remanei* and *C. latens* pure species males (n_males_=21 and n_spermatids_=506 for *C. remanei-latens* F1 hybrid males, n_males_=24 and n_spermatids_=645 for *C. remanei* pure species males, n_males_=26 and n_spermatids_=832 for *C. latens* pure species males; z=2.841, Tukey-corrected *P*<0.05; z=−2.964, Tukey-corrected *P*<0.05, respectively). **(C)** *C. remanei-latens* F1 hybrid males have sperm with lower average roundness (inverse-transformed) than *C. remanei* and *C. latens* pure species males (see sample sizes for spermatid circularity; z=3.328, Tukey-corrected *P*<0.01; z=−2.876, Tukey-corrected *P*<0.05, respectively). **(D)** *C. remanei-latens* F1 hybrid males showed a trend of fewer spermatids than *C. remanei* and *C. latens* pure species males or F1 male hybrids from the reciprocal cross (n=19 *C. remanei-latens* F1 hybrid males, n=20 *C. remanei* pure species males, n=20 *C. latens* pure species males, n=20 *C. latens-remanei* F1 hybrid males; z=2.278, *P*=0.0227; z=2.493, *P*=0.0127; z=2.104, *P*=0.0344 respectively), albeit not significant after multiple test correction (Tukey-corrected *P*>0.05 for all contrasts).

Spermatids isolated from 90 F1 males from conspecific and heterospecific crosses could be classified into three main categories of visible abnormalities: “spikey” spermatids, “separated” spermatids, and “budding” spermatids (Figure 3b-d, Table 3). Overall, more *C. remanei-latens* F1 hybrid males (33%) exhibited these morphological abnormalities compared to hybrid males from the reciprocal heterospecific cross (10%) and conspecific crosses (Table 3; 4% for F1 males from both *C. remanei* and *C. latens* conspecific crosses). For some individual males, most visible spermatids showed abnormal phenotypes, whereas other individuals had only a few abnormal spermatids at the time of imaging. In sum, in addition to defects in gonadogenesis and capacity to make spermatids at all, when they do produce sperm, *C. remanei-latens* F1 hybrid males tend to make phenotypically abnormal spermatid cells.

**Figure 3.**
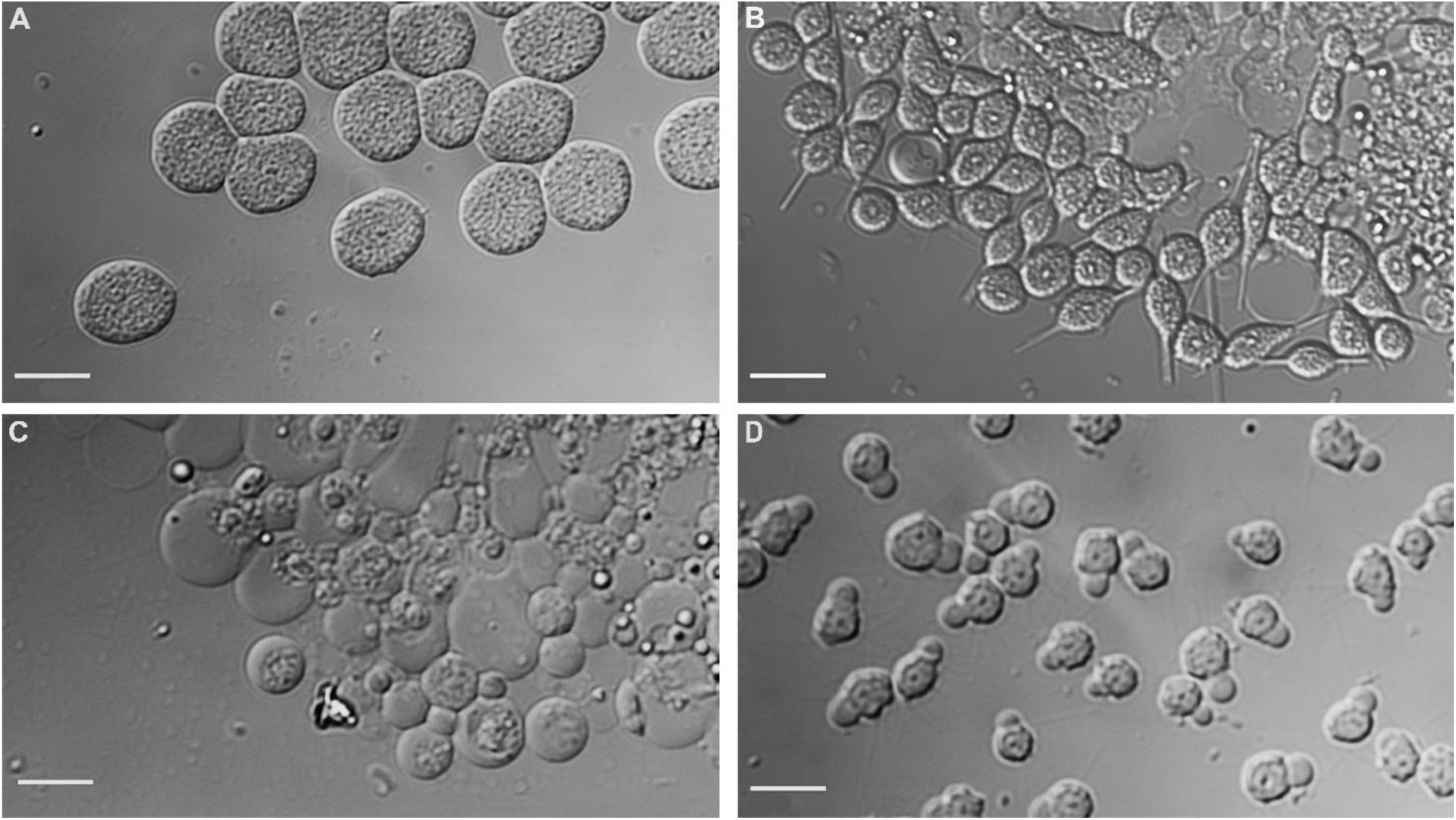
DIC images documenting variation in spermatid morphology. Scale bars represent 15µm. **(A)** Wild-type spermatids isolated from a *C. remanei* male. **(B)** Spikey spermatids isolated from a *C. remanei-latens* F1 hybrid male. **(C)** “Separated” spermatids isolated from a *C. remanei-latens* F1 hybrid male. **(D)** “Budding” spermatids isolated from an F1 male from a *C. latens-remanei* cross.

**Table 3.** Incidence of abnormal spermatid phenotypes in F1 male offspring from *C. latens* and *C. remanei* conspecific and heterospecific crosses.

| Parental cross* | n | # F1 males with normal spermatids | #F1 males with “spikey” spermatids | #F1 males with “separated” spermatids | #F1 males with “budding” spermatids |
| --- | --- | --- | --- | --- | --- |
| *C. remanei* x *C. remanei* | 24 | 23 | 1 | 0 | 0 |
| *C. remanei* x *C. latens* | 21 | 14 | 5 | 2 | 0 |
| *C. latens* x *C. remanei* | 20 | 18 | 0 | 1 | 1 |
| *C. latens* x *C. latens* | 26 | 25 | 1 | 0 | 0 |

\*Female x male

### Reciprocal cross asymmetry in hybrid tail malformation

Because disrupted gonadogenesis and spermatogenesis did not fully explain the patterns of hybrid male sterility individually, we investigated whether the male tail reproductive structures of hybrids also displayed abnormal phenotypes. Male tail structures are essential for mate apposition behavior as well as successful insemination (Koo et al. 2011), and any structural abnormalities could decrease an individual’s ability to reproduce. We found at least one structural tail defect in 40% of *C. remanei-latens* F1 hybrid males (n=125), an incidence significantly higher compared with *C. latens-remanei* F1 hybrid males (5.4%, z=−6.079, Tukey-corrected *P*<0.0001) and pure species F1 males (*C. remanei* parental: 6.7%, z=−5.098, Tukey-corrected P<0.0001; *C. latens* parental: 4.4%, z= −4.855, Tukey-corrected *P*<0.0001), all of which showed a similar likelihood of possessing at least one tail defect (Tukey-corrected *P*>0.05 for 3 pairwise comparisons) (Table 4). We detected no differences in the incidence of tail defects between pure species F1 males and *C. latens-remanei* F1 hybrid males (Table 4).

**Table 4.**
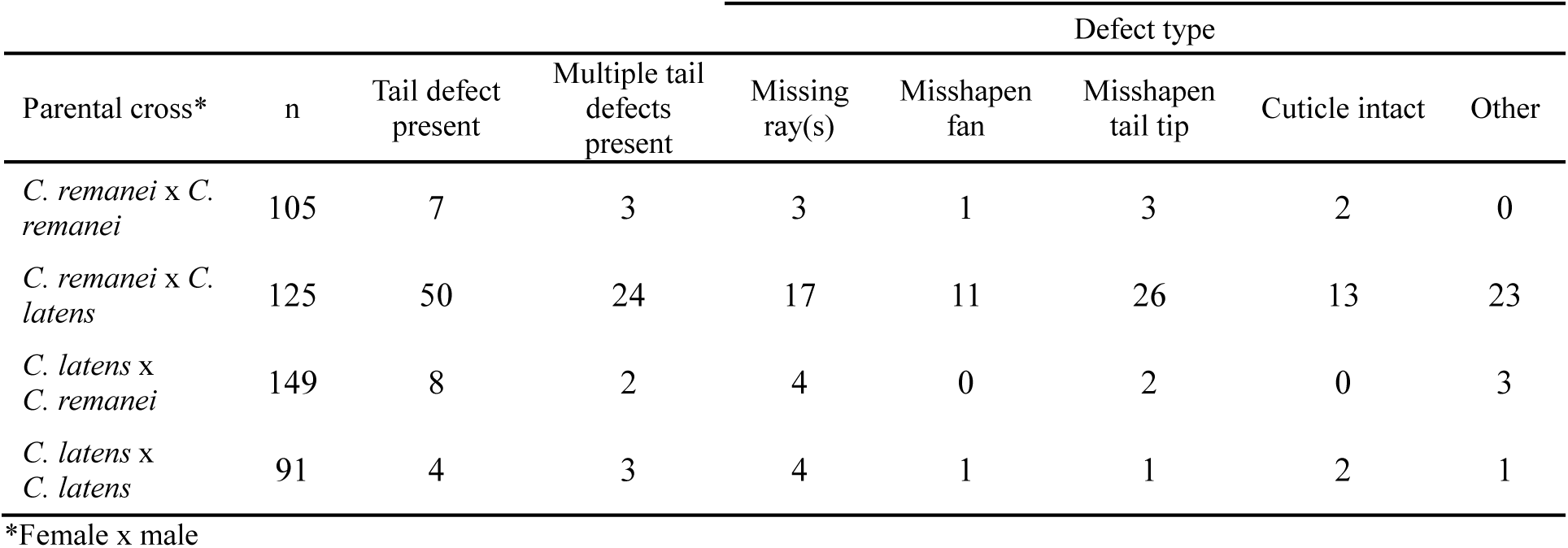
Number of F1 male offspring displaying abnormal tail morphology.

We scored each individual tail for the presence of 5 types of abnormalities: (i) missing or fused rays, (ii) abnormal fan shape, (iii) abnormal tail tip shape, (iv) failure to shed cuticle and (v) “other”. No abnormal spicule morphologies were detected among hybrid F1 males or conspecific F1 males. Abnormal *C. latens-remanei* F1 hybrid males (5.4% of n=149 F1 hybrid males) as well as *C. remanei* (6.7% of n=105 F1 males) and *C. latens* pure species F1 males (4% of n=91 F1 males) most often showed ray defects (43-100% of abnormal males from these cross types) (Table 4). Strikingly, however, 24 (19.2%) *C. remanei-latens* F1 hybrid males (n=125) displayed multiple abnormalities (Table 4), with 10% of individuals showing defects in 4 of the 5 categories. The most commonly malformed structure among *C. remanei-latens* hybrids was the tail tip (abnormal in 20.8% of individuals) followed by the rays (abnormal in 13.6% of individuals) (Table 4).

While malformed tail structures occur at a lower incidence than gonad abnormalities in *C. remanei-latens* F1 hybrid males, in combination with spermatid defects, they nonetheless contribute to a suite of morphological abnormalities associated with reproductive function that together result in the high degree of F1 hybrid male sterility in this cross direction.

## Discussion

### Postzygotic barriers between *C. remanei* and *C. latens* are multifarious

We investigated a diversity of reproductive traits in the nematodes *C. remanei* and *C. latens* and their hybrid male offspring, with special attention to F1 hybrid males produced from female *C. remanei* and male *C. latens* parents. Prior work has demonstrated such hybrid males suffer strong postzygotic reproductive isolation in the form of hybrid male sterility, which is not observed in the reciprocal cross direction (Dey et al. 2014; Bundus et al. 2018). We characterized a large collection of abnormal phenotypes in F1 hybrid males, including developmental defects in 1) the formation of somatic external reproductive structures of the male tail, 2) morphogenesis of the somatic gonad, and 3) gametogenic germline defects and abnormal spermatogenesis. This constellation of dysfunctional reproductive traits indicates that multiple developmental processes are disrupted in these male hybrids, all of which contribute to extensive overall hybrid male sterility.

#### Defects in F1 hybrid male somatic reproductive structures

We demonstrated that hybrid male tail defects are commonly associated with abnormal ray morphology and number. The innervated rays of the male tail are essential to initiate and maintain contact with females during mating, as well as to guide “searching” behaviors involved in vulva location (Koo et al. 2011). Missing or malformed rays can impede a male’s ability to perform these behaviors and lead to copulatory failure (Koo et al. 2011). Given the elevated frequency of under-developed, malformed, and missing rays among hybrid males, we predict that they experience difficulty performing mating behaviors and transferring sperm to females. Direct observation of mating activity could further evaluate the extent to which these tail irregularities prevent hybrid males from engaging in stereotypical mating behaviors to achieve successful intromission and ejaculation. However, only ∼40% of *C. remanei-latens* F1 hybrid males show tail defects, whereas >95% of them are sterile, indicating that defects to mating behavior and external reproductive morphology are insufficient to fully account for the observed magnitude of hybrid male sterility.

#### Abnormal gonad development in F1 hybrid males

An additional source of hybrid sterility for *C. remanei-latens* F1 hybrid males is malformation of gonad tissue itself (Dey et al. 2014). Disrupted development of gonadogenesis in hybrid males is common to *Drosophila* and other insects (Engels and Preston 1979; Goodpasture et al. 1980; Davis et al. 2020), mammals (Niayale et al. 2021), fish (Pinheiro et al. 2019), and other nematodes (Woodruff et al. 2010). In wild-type *Caenorhabditis* males, gonad development follows the J-shaped migration path of the linker cell during the larval stages, starting from a mid-ventral position of the linker cell that moves anteriorly then shifts to the dorsal side as it migrates posteriorly before shifting ventrally again toward the cloaca in the tail, finally undergoing programmed cell death just before adulthood to enable passage of sperm (Kimble and White 1981). Consistent with previous work (Dey et al. 2014), we showed that many gonad deformities of *C. remanei-latens* hybrid males implicate a failure to complete the expected migration path and/or connect to the cloaca. Early or incorrect termination of gonad elongation during development represents one possible explanation for our observation of excess spermatid accumulation and sterility in some of these males. In *C. elegans*, misexpression of *nhr-67* and other negative regulators of netrin receptors cause improper growth and positioning of the ventral gonad arm, but many genes involved in this period of gonad migration have not yet been characterized (Schwarz et al. 2012). Further studies that map genetic incompatibilities between *C. remanei* and *C. latens* will help to clarify which genes and pathways get disrupted in hybrids to cause disrupted hybrid male gonadogenesis.

Curiously, however, we found that the gonads of >40% of sterile *C. remanei-latens* hybrid males were nonetheless competent to make sperm, even when gonad tissue development was severely compromised. This observation indicates that gonad morphogenesis defects are partially independent of the ability for the germline to successfully perform spermatogenesis and, consequently, also is insufficient to fully explain the overall extent of hybrid male sterility. Although the hybrid males tended to produce fewer sperm cells when they were able to make them at all, in some individuals, we observed an excessive accumulation of sperm in the gonad and in several cases, somatic tissues. This excess sperm accumulation likely occurred due to a lack of connection of the gonad and the cloaca, consequently conferring an inability of the males to ejaculate. Similar accumulation of sperm has been observed in *C. elegans* mutants of *lin-27* and *let-7* due to linker cell death defects (Abraham et al. 2007); while premature linker cell death inhibits the completion of gonad elongation, it does not preclude germ cell proliferation (Kimble and White 1981). It is also possible that an abnormally long retention of spermatids could lead to precocious spermatid activation. Precocious sperm activation within the hybrid male gonad might lead to ectopic sperm cell migration, and analogous to the detrimental effect of ectopic sperm on females, corresponding tissue damage and reduced lifespan of the male. Consequently, sperm cell retention might act as a phenotypic link between hybrid male sterility and inviability, if accompanied by precocious activation, despite arguments that the genetic basis to compromised fertility and viability in hybrids generally ought to be decoupled (Presgraves and Meiklejohn 2021). Future experiments that test for differential survival in hybrid males with varying degrees of gonad and germline abnormalities can help elucidate this possible connection between hybrid male sterility and inviability.

#### Compromised spermatogenesis in F1 hybrid males

Given the ability of at least some hybrid males to make sperm, we examined their sperm cells and documented multiple atypical sperm cell phenotypes. Thus, even if mating and sperm transfer are successful, given the aforementioned incompleteness of those reproductive isolating barriers, sperm fertility may be compromised and contribute to hybrid male sterility. The sperm cell defects represent problems arising at different stages of spermatogenesis (meiotic production of haploid spermatids) and spermiogenesis (development of immature spermatids into activated mature spermatozoa competent for fertilization) (Ward et al. 1981; Kelleher et al. 2000). For example, we noted “spikey” cells in several *C. remanei*-*latens* F1 hybrid males that are reminiscent of *C. elegans* mutants of *fer-5* and *spe-15* that develop large crystalline inclusions after activation due to aggregations of major sperm protein (MSP) (Ward et al. 1981; Ward and Klass 1982; Kelleher et al. 2000). These inclusions inhibit proper pseudopod formation, structures that are necessary for sperm cell movement and contain proteins required to fertilize oocytes (Singson et al. 1998; Zannoni et al. 2003). Moreover, we found that spermatids derived from *C. remanei-latens* hybrid males were smaller and more irregularly shaped than spermatids of pure species individuals. Abnormal sperm development in hybrid males has clear implications for infertility, as defects in spermatogenesis have been documented in other taxa including mice and *Drosophila* (Dobzhansky 1934; Oka et al. 2010; Turner et al. 2012; Kanippayoor et al. 2020).

#### A constellation of dysfunctional hybrid phenotypes

Our analysis shows that hybrid male sterility arises due to the combined effects of dysfunctional traits that influence 1) copulation (e.g., due to abnormal tail reproductive structures or abnormal behavior resulting from improper tail neural development in the L3 or L4 stages), 2) transfer gametes (e.g., due to dysfunctional gonad morphogenesis or spermatogenesis), and 3) ability of sperm to fertilize oocytes (e.g., due to defective spermiogenesis or sperm cell migration behavior). Despite the influence of all three of these components, we find that disrupted gonad morphogenesis is the feature perturbed most prominently and profoundly among all the traits investigated. We also conclude that the L4 final larval stage appears to serve as a developmental hotspot for dysfunctional development in *C. remanei-latens* hybrid males: critical phases of gonad elongation and maturation, male tail morphogenesis, and early spermatogenesis all occur during this life stage (Emmons 2014). The dysfunctional development of these reproductive traits occur in addition to hybrid inviability due to embryonic arrest that disproportionately afflicts male hybrids (Dey et al. 2014; Bundus et al. 2018).

Our observations provide phenotypic support for the Fragile Male Hypothesis, which attributes accelerated hybrid male breakdown to the increased susceptibility of male-specific developmental programs to genetic perturbation (Wu et al. 1996; Cutter 2023, 2024). Faster molecular evolution of loci affecting male-specific development (Faster Male Hypothesis) or of loci on the X chromosome (Faster X Theory) also could, in part, explain why males specifically suffer increased developmental abnormalities (Orr and Presgraves 2000; Presgraves and Meiklejohn 2021; Cowell 2022; Cutter 2023, 2024). Further detailed analysis across multiple developmental stages in hybrids will help to determine whether earlier points in development mark crucial tipping points for phenotypic defects that fully manifest in L4 and adulthood (Cutter and Bundus 2020; Cutter 2023), both in *Caenorhabditis* specifically and generally across disparate taxa.

### Prezygotic barriers separating *C. remanei* and *C. latens* are undetectable

Prior to this study, *C. remanei* and *C. latens* heterospecific matings had been investigated only under “no-choice” conditions, where males of one species and females of the other species were exposed to one another (Dey et al. 2012, 2014; Bundus et al. 2018). We found that even in “choice” scenarios where both sexes of both species were present, *C. remanei* and *C. latens* males and females showed no consistent preference toward conspecific matings. Indiscriminate male-to-female and female-to-male long-distance attraction between species has been observed in more distantly related gonochoristic *Caenorhabditis* species due to shared responses to pheromone cues (Chasnov et al. 2007; Garcia et al. 2007; Borne et al. 2017), but the observed lack of positive assortative mating between *C. remanei* and *C. latens* is surprising given the incidence of extensive intrinsic postzygotic barriers that we observed. Nevertheless, these results suggest that strong premating barriers are not contributing to these species’ divergence.

Ectopic sperm cell migration serves as a prominent postmating-prezygotic barrier between certain *Caenorhabditis* species due to its detrimental effects on female fecundity and lifespan (Ting et al. 2018; Schalkowski et al. 2024). Although heterospecific crosses between *C. remanei* and *C. latens* produce significantly fewer offspring than do conspecific crosses (Dey et al. 2014), we observed ectopic sperm migration at similar frequencies following both types of matings. A detectable incidence of ectopic sperm after conspecific mating has been found in several other *Caenorhabditis* species, as well (Ting et al. 2014; Schalkowski et al. 2024). Low levels of ectopic sperm migration after conspecific mating may be an indicator of sexually antagonistic coevolution, potentially leading to arms-race coevolution between female traits and sperm traits that may get revealed in interspecies pairings (Rowe et al. 2003; Ting et al. 2018b; Cutter et al. 2019). Divergence in such a manner, however, does not appear to influence the outcome of heterospecific matings between *C. remanei* and *C. latens*. We therefore conclude that ectopic sperm migration does not represent a bona fide postmating-prezygotic barrier between *C. remanei* and *C. latens*.

### Drivers of divergence between *C. remanei* and *C. latens* remain unresolved

Sampling efforts to date suggest that *C. remanei* and *C. latens* occupy distinct geographic regions (Dey et al. 2012), although the degree to which human transportation has influenced their distribution is unknown (Cutter 2015; Frézal and Félix 2015). Many aspects of these species’ ecology also remain elusive, and we cannot yet say whether differences in abiotic microhabitat, microbial food source, or other biotic features could contribute ecological barriers. Although we are unable to draw conclusions about how ecological barriers could influence interspecies interactions in nature, if contact zones do exist, the lack of premating isolation suggests that ecological speciation in sympatry or parapatry is an unlikely driver of reproductive isolation between *C. remanei* and *C. latens*. We would expect early evolution of premating barriers because of divergent selection in differing environments if this were the case (Schluter 2001; Rundle and Nosil 2005), as natural selection tends to promote the fixation of traits that reduce competitive and reproductive exclusion between diverging groups (Schluter 2000; Pfennig and Pfennig 2009), but our results revealed strong intrinsic postzygotic isolation in benign laboratory conditions and no discernable premating barriers in this nutrient-rich environment. However, we cannot rule out the potential influence of extrinsic barriers in these species’ overall reproductive isolation, as empirical tests have used a common benign environment and tests of how distinct environmental factors differentially affect their reproductive isolation have not yet been conducted. Based on available evidence, we suggest that *C. remanei* and *C. latens* have primarily diverged in allopatry and hypothesize that they have not been subject to substantial reproductive or ecological character displacement. Such circumstances would be consistent with intrinsic isolation accumulating as in theories of “systems drift”, which can generate potent genetic incompatibility on reasonable time scales even through neutral accumulation of changes in gene regulatory networks (GRNs) (Schiffman and Ralph 2022). The geographic separation of most known isolates of *C. remanei* and *C. latens* also raises the prospect that further sampling may discover additional closely related species with partial reproductive isolation that could aid in further testing for extrinsic barriers and premating barriers on even shorter timescales of divergence.

### Many DMI loci or few DMI loci with complex pleiotropic effects?

*C. remanei*-*latens* F1 hybrids show clear signatures of intrinsic postzygotic isolation involving multiple disrupted developmental processes. However, it’s unclear whether the disparate dysfunctional reproductive phenotypes arise through shared or distinct genetic pathways. Numerous studies in other organisms have uncovered a polygenic basis for hybrid sterility and inviability (Reed et al. 2008; Arnegard et al. 2014; Thompson et al. 2022), including other species of *Caenorhabditis* (Bi et al. 2015, 2019). However, genetic mapping of DMIs between *C. remanei* and *C. latens* is thus far limited (Bundus et al. 2018), making it difficult to truly pinpoint the number of discrete elements contributing to any given barrier trait.

Previous work proposed that hybrid sterility in *C. remanei*-*latens* F1 hybrid males may be underpinned by a simple X-autosome DMI whereas hybrid inviability requires explanation by many incompatible loci (Bundus et al. 2018). Our observations, however, reveal the capacity of hybrids to show multiple distinct sterility-causing phenotypes, suggesting a genetic architecture that might be more complex than previously appreciated. The greater complexity could manifest in multiple possible ways that, at present, we are unable to distinguish. First, to be consistent with backcross analyses (Bundus et al. 2018), one or just a few X-autosome DMIs might occur to cause hybrid male sterility but arise from the polygenic interaction of many contributing genes (Satokangas et al. 2020). Second, one or just a few DMIs might occur but lead to a diversity of distinct pleiotropic phenotypic effects. For example, inter-species introgression studies of *C. briggsae* and *C. nigoni* show that a given introgressed chromosomal region can lead to strong hybrid male sterility as well as hybrid inviability (Bi et al. 2019). Third, the genetic architecture of dysfunctional male traits might involve many distinct DMIs, albeit most with modest effect on overall sterility despite quantifiable phenotypes, such that previous backcross analysis of hybrid male sterility underestimated the numerical extent of genetic incompatibilities. In the absence of high-resolution genetic mapping, we cannot distinguish these alternatives, and further work that genetically maps and characterizes incompatibility loci will help determine the roles of multiple loci and pleiotropy in generating dysfunctional phenotypes of hybrids.

Interestingly, not all hybrid males show the same combination of developmental abnormalities. Even among sterile hybrids, we observe variation in which phenotype imposes the strongest barrier to reproduction. Some individuals fail to complete gonad morphogenesis and thus cannot release sperm through their cloaca, while others have relatively normal gonads but fail to produce sperm at all. Whether these phenotypes are the product of separate DMIs, the result of incomplete penetrance of incompatibilities, or are subject to epigenetic modification remains to be determined, but it is clear that a complex developmental genetic architecture governs *C. remanei*-*latens* F1 hybrid male sterility.

All together, our findings indicate a constellation of reproductive barriers that contribute to intrinsic postzygotic isolation between *C. remanei* and *C. latens* in their male hybrid offspring in a manner consistent with Haldane’s rule (and Darwin’s corollary to Haldane’s rule). Our observations are consistent with the fragile male and faster male hypotheses and highlight the L4 stage as a hotspot for developmental irregularities. We find no evidence for strong premating or postmating-prezygotic isolation between *C. remanei* and *C. latens* but describe diverse developmental consequences of postzygotic genetic incompatibilities by characterizing a suite of traits associated with hybrid male sterility. We conclude that postzygotic reproductive isolation evolves faster than prezygotic isolation in this system, framing *Caenorhabditis* as an intriguing contrast to other organisms that exhibit a reciprocal temporal pattern of accumulation of reproductive isolating barriers.

## Author contributions

M.N.D. carried out the assortative mating, ectopic sperm migration and hybrid tail defect experiments and completed preparation of the initial article draft. A.P. carried out the hybrid fertility, spermatid morphology and spermatid defect experiments, and D.I. participated in conceptualization and data collection for these experiments. J.B.D.S. participated in conceptualization of the project and performed informative pilot experiments prior to the study. A.D.C. edited and reviewed article drafts, acquired funding for the project and provided supervision and guidance during the planning and execution of laboratory experiments.

## Data availability statement

The data underlying this article are available through Zenodo at https://dx.doi.org/10.5281/zenodo.14888936 and on GitHub at https://github.com/Cutterlab/Constellation_of_RIs.

## Acknowledgements

We are grateful to Rebecca Schalkowski for advice on performing experiments and feedback on the manuscript. D.I. was supported by an Undergraduate Summer Research Award from the Natural Sciences and Engineering Research Council of Canada (NSERC USRA). This work was also supported by a Discovery Grant to A.D.C. from the Natural Sciences and Engineering Research Council of Canada.

## Supplementary figures

**Supplementary figure 1.**
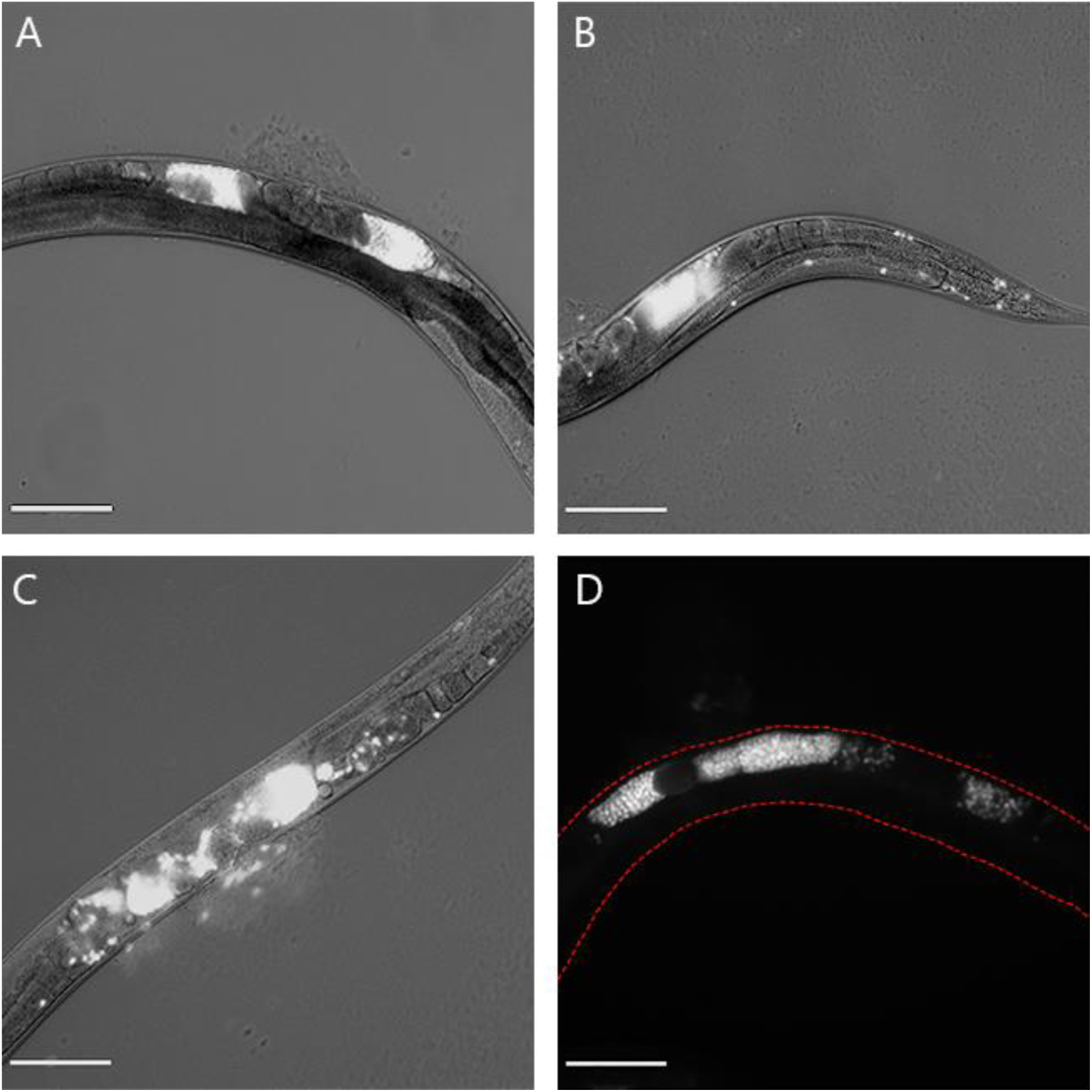
Images of fluorescent sperm within *C. remanei* and *C. latens* females after mating with males dyed with MitoTracker Red CMXRos. Bright areas indicate presence of fluorescent sperm. Scale bars represent 50µm. (A) *C. latens* female containing the sperm of a *C. remanei* male. Sperm are restricted to the uterus. (B) *C. latens* female displaying mild ectopic sperm migration in gonadal and somatic tissues after mating with a *C. remanei* male. (C) *C. remanei* female showing moderately severe ectopic sperm migration in gonadal tissue after mating with a *C. latens* male. (D) *C. remanei* female showing severe ectopic sperm migration in gonadal tissue after mating with a *C. latens* male.

